# Time to fixation in changing environments

**DOI:** 10.1101/2021.05.04.442623

**Authors:** Sachin Kaushik, Kavita Jain

## Abstract

Although a large number of experimental and theoretical studies have been carried out in a constant environment, as natural environments vary in time, it is important to ask if and how these results are affected by a changing environment. Here, we study the properties of the conditional fixation time of a mutant in a finite, randomly mating diploid population which is evolving in a periodically changing environment. In a static environment, as the conditional mean fixation time of a co-dominant beneficial mutant is equal to that of a deleterious mutant with the same magnitude of selection coefficient, similar patterns for beneficial and deleterious sweeps may result. We find that this symmetry breaks even when the environment is changing slowly. Furthermore, for intermediate dominance, the conditional mean fixation time of a beneficial mutant in a slowly changing environment depends weakly on the dominance coefficient and is close to the corresponding results in the static environment; however, the fixation time for a deleterious mutant under moderate selection with a slowly varying selection coefficient differs substantially from that in the constant environment when the mutant is recessive. Our results thus suggest that the variability patterns and levels for beneficial sweeps are mildly affected by temporally varying environment but changing environment is likely to strongly impact those due to recessive deleterious sweeps.

## 1 Introduction

In a finite, recombining population where a beneficial locus is linked to neutral loci, if the advantageous mutation fixes sooner than the time it takes to get dissociated from the linked loci via recombination, the neutral genetic diversity in the neighborhood of the selected locus is reduced (beneficial sweeps) (Maynard Smith and Haigh, 1974; Stephan, 2016); similarly, deleterious sweeps, mainly, due to the fixation of mildly deleterious mutations can also occur. Thus, the time of fixation is intimately related to the level and patterns of neutral diversity (Tajima, 1990).

Theoretical models of sweeps and their genomic applications assume the environment to be constant in time; however, natural environments vary which can affect the fixation time. For example, if a mutant arises when the selection is positive and increasing, the mean fixation time, conditional on fixation, is expected to be smaller than when the selection pressure remains the same as that when the mutant arose, thus leading to a more reduction in the neutral diversity in the changing environment. One may then ask: how much does the fixation time in a changing environment (especially, if it varies slowly) differ from that in the static environment?

Furthermore, in static environments, an important property of the conditional mean fixation time is that it is same for a mutant with selection coefficient *s* and dominance coefficient *h* and a mutant with respective parameters, −*s* and 1 − *h* (Maruyama, 1974; Maruyama and Kimura, 1974), as a result of which it may be difficult to distinguish between the diversity patterns due to positive and negative selection (Johri *et al*., 2020). In a changing environment, on general grounds, this symmetry can be expected to break, and one may then enquire about the extent to which this symmetry is broken and delineate the parameter space where it is strongly broken.

As a first step towards an understanding of sweeps in changing environments, here we study the properties of conditional fixation time of a mutant in a finite, diploid population when the selection coefficient is time-dependent. To the best of our knowledge, except for a preliminary study (Uecker and Hermisson, 2011), these have not been investigated in detail. We consider the evolution in an environment that changes periodically due to, for example, seasonal cycles (Williams *et al*., 2017), and study how the fixation time is affected by the rate of environmental change, time of appearance of the mutant, strength of selection and the dominance coefficient. Throughout the article, we assume random mating and autosomal inheritance.

Our results are obtained analytically using a diffusion theory for time-inhomogeneous processes when the mutant with either sign of selection is under weak or moderate selection and a semi-deterministic theory for strongly selected beneficial mutants, and are supplemented and checked by numerical simulations. Our main finding is that in slowly changing environments, the conditional mean fixation time can be well-approximated by that in the static environment for beneficial mutants with intermediate dominance and deleterious dominant mutants; however, this result does not hold for recessive deleterious mutants under moderate selection. It then follows that the symmetry property for conditional mean fixation time mentioned above (Maruyama, 1974; Maruyama and Kimura, 1974) is strongly broken between the recessive deleterious and dominant beneficial mutants. Since by virtue of Haldane’s sieve that operates in both static (Haldane, 1927) and slowly varying environments (Devi and Jain, 2020), most deleterious mutations are recessive and beneficial ones are dominant, our results can be relevant to an understanding of selective sweeps.

## 2 Model

We consider the model in Devi and Jain (2020) that deals with a randomly mating population of size *N* . We assume that a single biallelic locus is under selection and the three genotypes, *aa, Aa* and *AA* have the fitness 1 + *s*(*t*), 1 + *hs*(*t*) and 1, respectively. Here 0 *< h <* 1 is the dominance parameter and 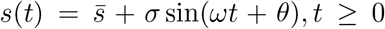 is the time-dependent selection coefficient that varies periodically with cycling frequency *ω*. Without loss of generality, we assume that the oscillation amplitude *σ >* 0 but the time-averaged selection coefficient 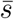 is arbitrary, and the initial phase 0 *≤ θ <* 2*π*.

For large populations and weak selection, instead of genotypic frequencies, we can work with the frequency *p* of the mutant allele *a* and *q* = 1 − *p* of the wildtype allele *A* (Nagylaki, 1992). We start with a single mutant allele in the population and ignore any further mutations. The evolution of the population under selection and random genetic drift is modeled by a continuous time birth-death process (Chapter 3, Lawler (2006)) in which the number *i* of allele *a* increases or decreases by one at rate *r*_*b*_(*t*) or *r*_*d*_(*t*), respectively, that are given by

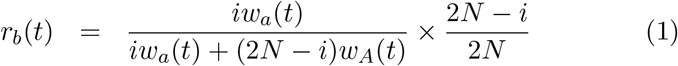

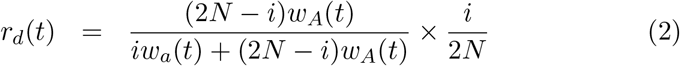

where *w*_*a*_(*t*) = (1 + *s*(*t*))*i* +(1 + *hs*(*t*))(2*N* − *i*) and *w*_*A*_(*t*) = (2*N* − *i*) + (1+*hs*(*t*))*i* are, respectively, the marginal fitness of allele *a* and *A*. The allele numbers at time *t* are updated at time *t* + *δt* where the interval *δt* follows an exponential distribution with rate, 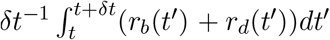.

For computational efficiency, numerical simulations of the above model were carried out assuming that the birth and death rates remain constant during the interval *δt*; however, we have checked that our results do not change if we relax this assumption. The simulation results for fixation time were obtained by averaging over 10^3^ and 10^4^ fixation events for deleterious and beneficial mutants, respectively.

## 3 Symmetry-breaking between conditional fixation times

In a constant environment, diffusion theory predicts that the mean fixation time of a mutant (*p →* 0), conditioned on fixation has a remarkable property: it is same for a beneficial mutant with selective advantage *s* and dominance coefficient *h* and a deleterious mutant with respective parameters −*s* and 1 − *h* (Maruyama, 1974; Maruyama and Kimura, 1974). Furthermore, for a codominant mutant (*h* = 1*/*2), this symmetry holds for any 0 *< p <* 1 (p. 170, Ewens (2004)).

In order to test whether the above result holds in a periodically changing environment, we need to consider mutants whose selection coefficient are of opposite sign at *all* times. Figure 1 shows the conditional mean fixation time 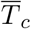 for mutants with dominance parameter *h* and 1 − *h* when they are neutral on-average but have selection coefficient *s*(*t*) and −*s*(*t*), respectively (for nonzero 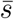, see Fig. S1). At low cycling frequencies (*ω* ≪ *N* ^−1^, *σ*), as expected by continuity, the time 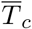 for both mutants is close to that in the static environment but the aforementioned symmetry is broken. At high cycling frequencies (*ω ≫ N* ^−1^, *σ*), as the population is sensitive to the time-averaged selection coefficient, the fixation time for both mutants approaches the neutral expectation, namely, 2*N* (Kimura, 1957) with increasing *ω* but the two fixation times do not coincide with each other. At intermediate frequencies, the conditional mean fixation times between the two mutants differ substantially and exhibit an extremum at the resonance frequency *ω*_*r*_ which is proportional to the inverse population size when the time-averageselection coefficient 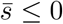 and 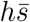 for 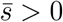 (Devi and Jain, 2020).

**Figure 1:**
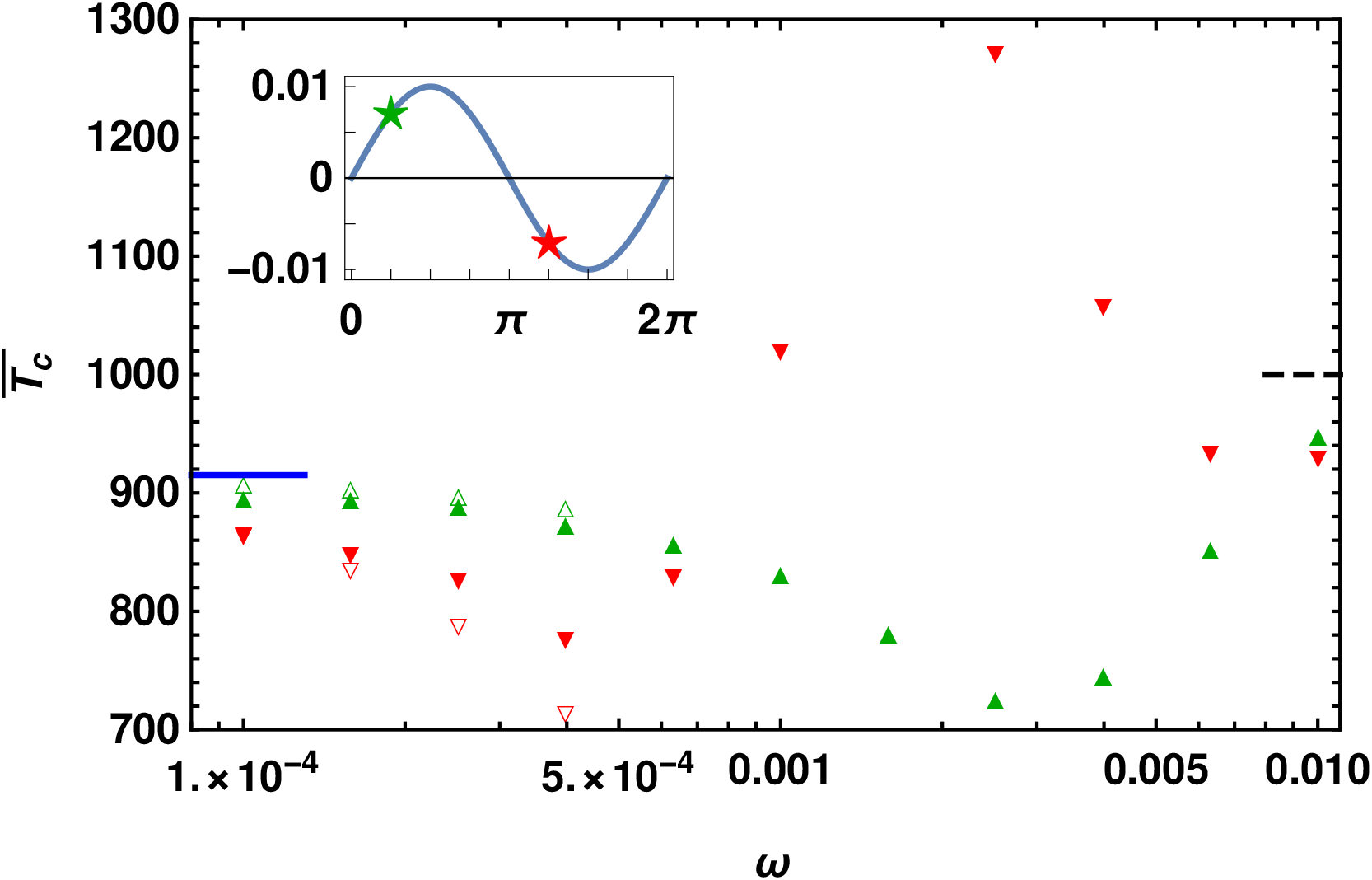
Conditional mean fixation time 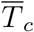 of a dominant mutant (*h* = 0.7, 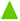) and a recessive one (*h* = 0.3, ▾) with selection coefficient *s*(*t*) = *σ* sin(*ωt* + *π/*4) and −*s*(*t*), respectively (see inset for *s*(0)) to show that the symmetry between the conditional mean fixation time for dominant beneficial and recessive deleterious mutant in static environments is not preserved in changing environments. The data is obtained by numerical simulations (closed symbols) and numerically integrating the diffusion theory equations (3) and (4) (open symbols) for small cycling frequencies. The conditional mean fixation time in the static environment with selection |*s*(0) | (solid line) and in the neutral environment given by 2*N* (dashed line) are obtained from diffusion theory and shown for comparison. In all the cases, *σ* = 0.01 and *N* = 500 so that *α* = *N* |*s*(0)| ≈ 3.5.

### 3.1 Fixation time in slowly changing environments

Importantly, Fig. 1 shows that in slowly changing environments, although the conditional mean fixation time for the beneficial mutant (for which selection remains positive until it fixes) deviates by a small amount from the corresponding result in the static environment with selection coefficient *s*(0), the fixation time for the deleterious mutant differs substantially from the constant-environment result.

To explore and understand this observation, we develop a diffusion theory with time-dependent selection coefficient in Appendix A. In periodically changing environments, simple expressions for the eventual fixation probability have been obtained in Devi and Jain (2020) for slow and fast changing environments using a perturbation theory. Below, using the same method, we find the mean fixation time in slowly changing environments. Writing the unconditional mean fixation time and eventual fixation probability as 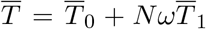 and *u* = *u*_0_ + *Nωu*_1_, respectively, in (A.4) and (A.5) and matching terms of the same order in *Nω* on both sides of the equations, we find that *u*_1_ and 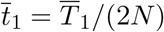 that capture the effect of changing environment obey the following ordinary differential equations,

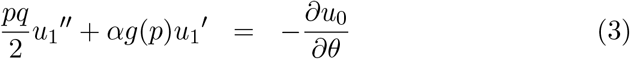

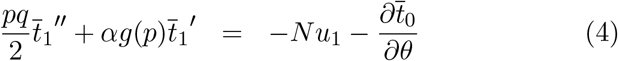

where the quantities *u*_0_ and 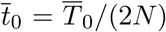 in the static environment obey (Kimura and Ohta, 1969)

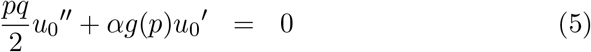

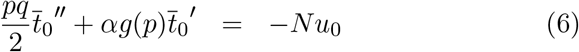

In the above equations, *g*(*p*) = *pq*(*p* + *h*(1 − 2*p*)) and *α* = *Ns*(0) is the scaled selection strength. Equations (3)-(6) are subject to the boundary conditions *u*_0_(0) = *u*_1_(0) = *u*_1_(1) = 0, *u*_0_(1) = 1 and 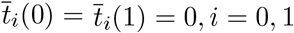 To linear order in *Nω*, the conditional mean fixation time is then given by

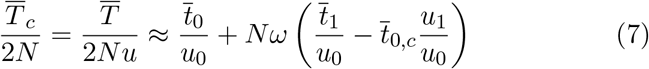

As attested by Fig. 1, the results obtained by numerically integrating (3)-(6) are in good agreement with those from simulations for small cycling frequencies. Although a formal solution to (7) can be written down, as it involves multi-dimensional integrals, it appears difficult to obtain a simple analytical expression for arbitrary *h* and 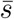. In the following, we will therefore study the weak, moderate and strong selection regimes separately using appropriate techniques.

### 3.2 Selective strolls in slowly changing environments

In a recent numerical study, it was found that in a static environment, for a mutant under weak selection (*N* |*s*| ≪ 1) and with intermediate dominance, the conditional mean fixation time *increases* with selection pressure and can be larger than that for a neutral mutant (Mafessoni and Lachmann, 2015). To see this result, for small *α* = *Ns*, we expand *u*_0_ and 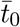 in a power series about *α* = 0 to order *α*^2^ and substitute them in (5) and (6). Matching terms of the same order in *α* on both sides of the equations, we get a set of second order differential equations which can be solved straightforwardly. For *p →* 0, we finally obtain

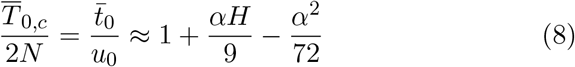

where *H* = *h* − (1*/*2) is the deviation from codominance. The above result shows that the conditional mean fixation time (relative to the neutral fixation time) is a non-monotonic function of *α* with a maximum at *α*^*\**^ = 4*H* and the corresponding time 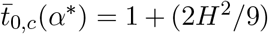. As *H*^2^ *<* 1*/*4, the conditional fixation time of the selected mutant can exceed that of the neutral mutant at most by *∼* 5%, as observed numerically in Mafessoni and Lachmann (2015).

In slowly changing environments that are neutral on-average 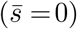, we proceed in a similar fashion as that for static environments; that is, we expand *u*_1_(*p*) and 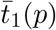 in a power series in *α* to quadratic order and plug them in (3) and (4). We then find

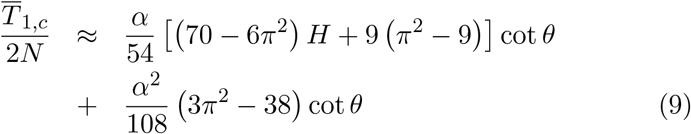

Then the maximum in the total conditional mean fixation time occurs at

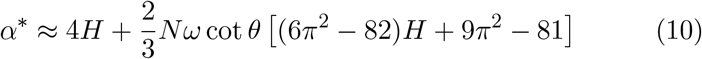

Equations (8) and (9) as also Fig. 2 show that the conditional mean fixation time continues to be a non-monotonic function of selection strength in changing environments. But for a codominant mutant, while 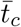 is symmetric about *α* = 0 in constant environment, due to symmetry-breaking in the changing environment, the maximum in the fixation time occurs at a negative *α* if the mutant arises when the selection is decreasing towards 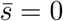 and positive *α* otherwise, as predicted by (10). For small *α*, from (9), we have 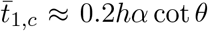 which shows that the changing environment has the strongest effect when the mutant is dominant; however, the magnitude of these effects is quite small even in these extreme cases, see inset of Fig. 2.

**Figure 2:**
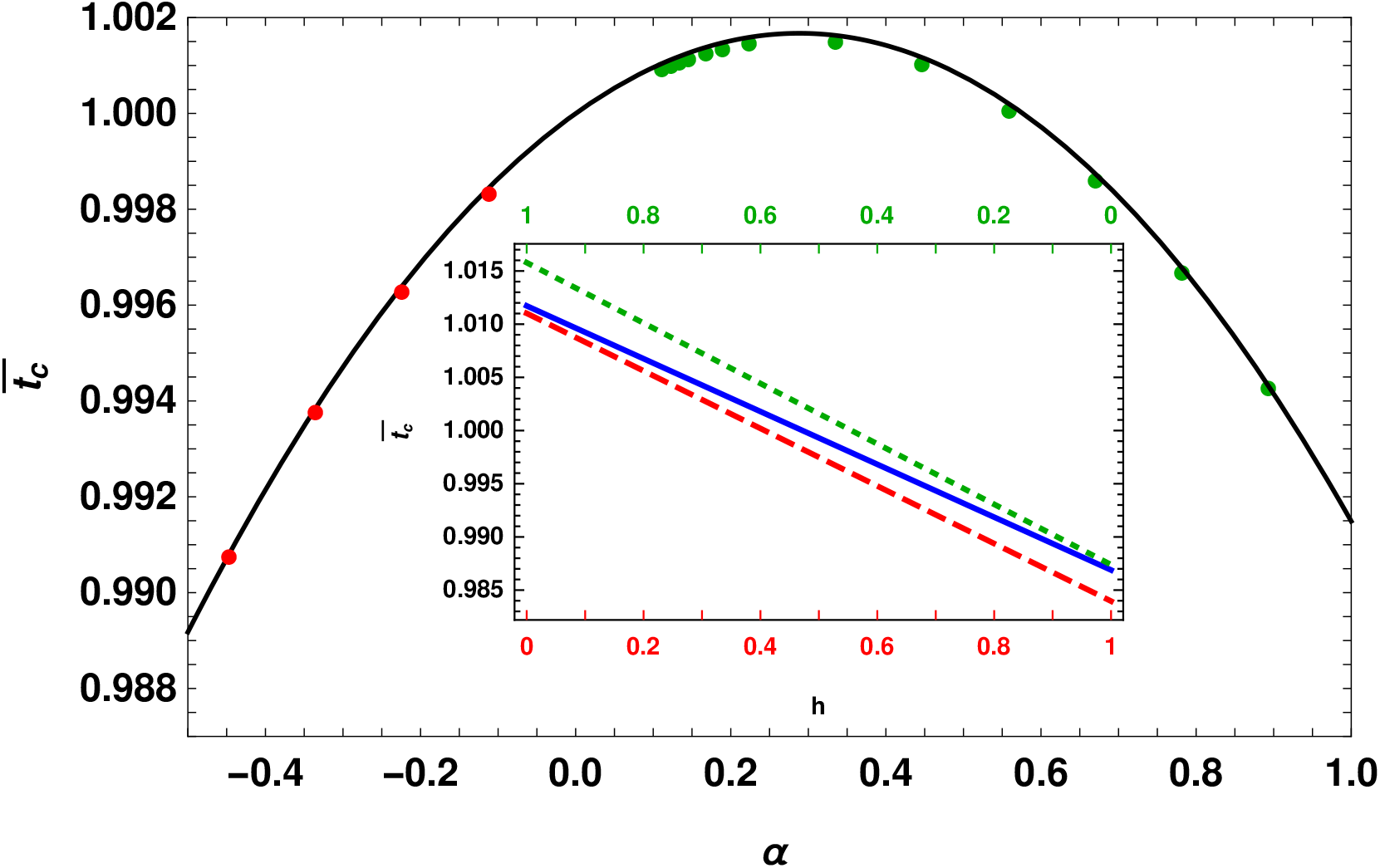
Scaled conditional mean fixation time, 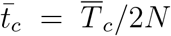 for a co-dominant mutant under weak selection in slowly changing, neutral on-average environment. The parameter *α* was varied with *σ*, keeping *N* and *θ* = *π/*4 (for positive *α*) and 5*π/*4 (for negative *α*) fixed. The inset shows the variation of conditional mean fixation time with dominance in static environment (solid) and slowly varying environment for initial phase *θ* = *π/*4 (dotted) and 5*π/*4 (dashed), and *σ* = 0.000158. In both plots, *N* = 2 × 10^3^ and *Nω* = 0.08, and the lines show the analytical expressions (8) and (9) and points shows the numerical solution of (3)-(6).

### 3.3 Deleterious sweeps in slowly changing environments

We now consider the parameter regime where selection is moderately strong (1 ≪ *α* ≲20). As in the last subsection, one would like to obtain simple analytical expressions for the time 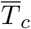 but, unfortunately, it is generally not possible to develop consistent approximations when the parameters are of moderate size. Below, we therefore rely on numerical solution of (3)-(6) and semi-quantitative arguments.

In static environments, for a mutant under moderately strong, positive selection, as the recessive mutant has a smaller probability to fix than the dominant mutant (Haldane, 1927), it must establish sooner to escape stochastic loss (Teshima and Przeworski, 2006); similarly, for negative selection, as the dominant mutant is more likely to get lost, its conditional fixation time is expected to be shorter than that for the recessive mutant. As Fig. 3 shows for |*α*| ≈ 14, these trend continue to hold in slowly changing environments as well. Also, as in Fig. 1, we find that the conditional mean fixation time for the beneficial mutant is quite close to the corresponding result in static environment for all dominance values shown. But for negative selection, the recessive mutant in changing environment, depending on the initial phase, takes far longer or shorter than the mutant in constant environment with selection coefficient *s*(0).

**Figure 3:**
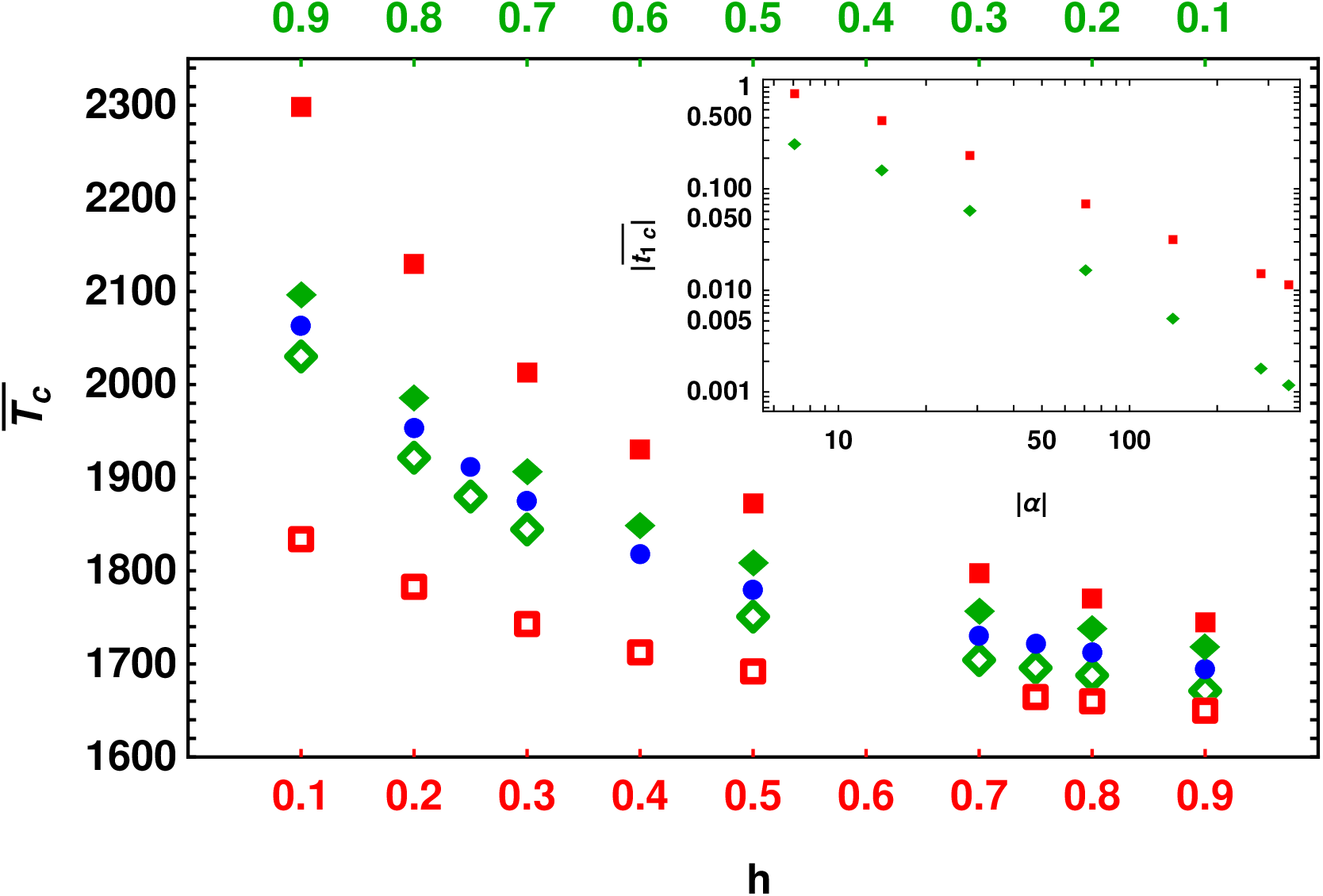
Conditional mean fixation time for moderate selection in static (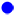) and slowly changing environment with selection coefficient *s*(*t*) = *σ* sin(*ωt*+*θ*) (diamonds) and −*s*(*t*) (squares) for different dominance coefficients and the initial phase *θ* = *π/*4 (open symbols) and 3*π/*4 (closed symbols). The other parameters are *N* = 2 × 10^3^, *Nω* = 0.05 and *σ* = 0.01. The inset shows the deviation between conditional mean fixation times in slowly changing and static environments for beneficial (*θ* = *π/*4, 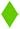) and deleterious mutants (*θ* = 5*π/*4, 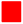) as a function of |*α* | for fixed *h* = 1*/*2 and *σ* = 0.01. In both plots, the data are obtained within the framework of diffusion theory by numerically solving (3)-(6).

To rationalize the qualitative behavior described above, we first note that due to (4) and (7), the time 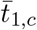 that captures the effect of slowly changing environment on conditional fixation time depends on the change *u*_1_ in the fixation probability which was studied recently (Devi and Jain, 2020). From (11b) of Devi and Jain (2020) for a beneficial mutant in a slowly changing, on-average neutral environment, we find that the change in the fixation probability due to changing environment is independent of *h* which suggests that the change in the fixation time is also same for any dominance coefficient. This argument is consistent with the data in Fig. 3. In contrast, for the deleterious mutant, (11c) of Devi and Jain (2020) shows that the change in the fixation probability is proportional to (1 − *h*)*/h* so that the fixation time in varying environment differs significantly from that in constant environment for recessive mutants, as seen in Fig. 3.

As in static environments, the conditional mean fixation time decreases with increasing selection strength in changing environments also. The inset of Fig. 3 shows that for large |*α*|, the correction in fixation time due to the changing environment decays faster for the beneficial mutant than the deleterious one. For strong selection, in the following section, we will show that 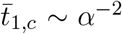 when the mutant is beneficial at all times and in Appendix B, we find that the corresponding correction for a deleterious mutant decays much slowly, as |*α*|^−1^; we thus conclude that the changing environment has a much stronger impact on deleterious mutations.

The conditional mean fixation time also depends on the initial phase of the selection coefficient. If the mutant arises when the magnitude of selection is decreasing towards 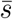, it fixes more slowly than in the static environment while the fixation time is less if the mutant arises when the selection is increasing in time. These qualitative behavior are consistent with those in the static environment where the fixation time increases with decreasing magnitude of selection pressure.

## 4 Fixation time of a beneficial mutant in large population

We now turn to the regime where selection is strong (|*α*| *≫* 10^2^). As the chance of fixation of a deleterious mutant is negligible for such strong selection (Kimura, 1957), here we focus on the fixation time of a mutant with positive time-averaged selection coefficient 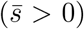, and study its dependence on the dominance coefficient and rate of environmental change within a semi-deterministic theory (Cohn and Jagers, 1994). This approach was recently used to find the distribution of conditional fixation time in a constant environment (Martin and Lambert, 2015); here, we are interested in generalizing these results to time-dependent environments.

Starting at a low initial frequency, if it escapes stochastic loss, the mutant population evolves stochastically until a time *t*_1_ when it reaches a finite frequency (phase *A*). For such trajectories, it is a good approximation to treat the further evolution of the mutant population deterministically (phase *B*). However, at a time *t*_2_(*> t*_1_), when the mutant frequency is close to one, as the wildtypes are in low numbers, they are subject to stochastic fluctuations and go extinct at a time *T*_*c*_ (phase *C*). The stochastic phases *A* and *C* can be described using a Feller process, as discussed below.

### 4.1 Time-inhomogeneous Feller process

In a time-dependent environment, the mutant allele frequency distribution Φ_f_(*p, t*|*p*_0_, 0) obeys the following forward Kolmogorov equation (Risken, 1996)

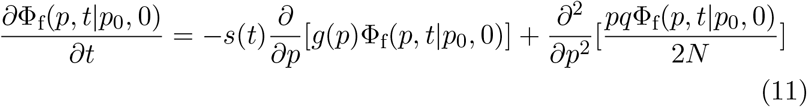

where, as before, *g*(*p*) = *pq*(*p* + *h*(1 − 2*p*)) and 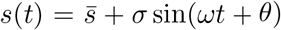. At short times where the mutant frequency is low (*p →* 0), the frequency distribution Φ_f_ *→ℱ* and (11) reduces to

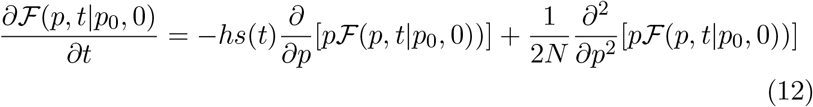

where *ℱ* (*p, t*|*p*_0_, 0) is the probability distribution of a Feller process (Feller, 1951a,b). (We will use the Feller process to describe the wildtype dynamics also at large times where the wildtype frequency is low.) Besides the evolutionary dynamics considered here, Feller diffusion has been used in modeling various other biological and physical processes (Cattiaux *et al*., 2009; Masoliver and PerellÓ, 2012; Gan and Waxman, 2015; Masoliver, 2016), and can be easily generalized to include mutations and time-dependent population size.

The exact probability distribution *ℱ* (*p, t*) of the mutant frequency with the initial condition *ℱ* (*p*, 0) = *δ*(*p* − *p*_0_) is given by (see Appendix C),

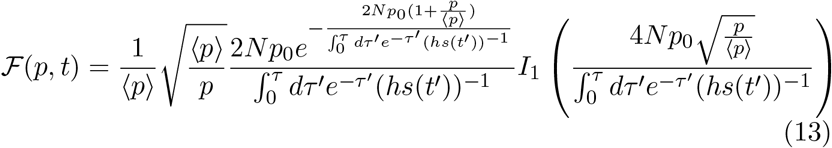

where *I*_*n*_(*z*) is modified Bessel function of the first kind, and

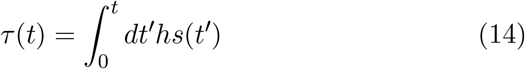

In (13), 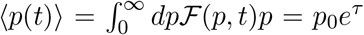 is the expected mutant allele frequency at time *t*, as can also be checked using (12).

### 4.2 Fixation probability

Since the probability that the mutant dies out by time *t* is equal to 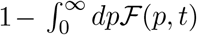, its eventual fixation probability, 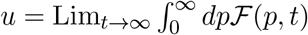 (see also Appendix C); using (13) and (14), we then obtain

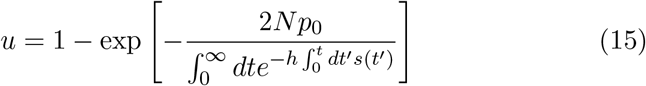

The above equation shows that *u* is nonzero provided the integral 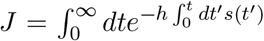 is finite. We verify that for constant selection and single initial mutant, (15) gives *u* = 1 − *e*^−*hs*^ ≈ *hs, h >* 0 for a beneficial mutant (Haldane, 1927). For time-dependent selection, the fixation process is time-inhomogeneous which means that *u* is a function of the initial phase *θ*.

Before proceeding further, we compare the result (15) with that obtained using a birth-death process in earlier studies (Kendall, 1948; Uecker and Hermisson, 2011; Devi and Jain, 2020). While *u*^(Feller)^ = 1 − *e*^−1*/J*^, the probability *u*^(birth-death)^ = (1 + *J*)^−1^ (refer (4) of Devi and Jain (2020)) for single mutant. For small selection coefficients, as the fixation probability is expected to be small, *J* must be large. Then, it follows that to leading order in 1*/J*, both the processes yield the fixation probability to be 1*/J* . For later reference, we recall that the fixation probability in a periodically changing environment exhibits an extremum at the resonance frequency *ω*_*r*_ which is proportional to the growth rate 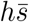 of the mutant (Devi and Jain, 2020).

### 4.3 Distribution of fixation time

In the stochastic phase *A*, although the expected mutant frequency grows exponentially with time, (13) and (15) show that at large times (where *τ → ∞*, since *s*(*t*) *>* 0 at all times), the random variable *y* = *p/*⟨*p*⟩, conditioned on fixation, has a stationary distribution,

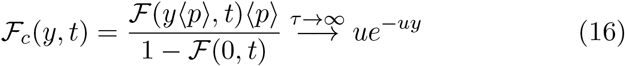

(see also Uecker and Hermisson (2011)). Furthermore, in the vicinity of time *t*_1_, the mutant frequency in the stochastic phase *A* is given by *p* = (*p*_0_*y*)*e*^*τ*(*t*)^, *t ≲ t*_1_ and in the deterministic phase *B*, the average mutant frequency grows as 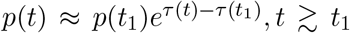. Thus the initial mutant frequency in the deterministic phase is given by 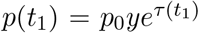, and from (16), it follows that the random variable 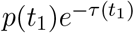 is exponentially distributed with mean (2*Nu*)^−1^.

In the deterministic phase *B* that begins at time *t*_1_ and ends at time *t*_2_, the average mutant frequency obeys *dp/dt* = *s*(*t*)*g*(*p*); integrating this equation over time from *t*_1_ to *t*_2_, we get *τ* (*t*_2_) − *τ* (*t*_1_) = *h*[*𝒟* (*p*(*t*_2_)) −*𝒟* (*p*(*t*_1_))], where

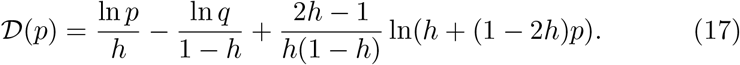

But as the frequency *p*(*t*_1_) *→* 0, *q*(*t*_2_) *→* 0, the initial frequency *q*(*t*_2_) in phase *C* is related to *p*(*t*_1_) as

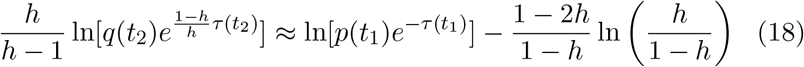

In the stochastic phase *C*, the wildtype population evolves stochastically from time *t*_2_ until it goes extinct at time *T*_*c*_. The wildtype frequency can be described by a Feller process that obeys (11) for the distribution 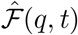 when *p → q, s*(*t*) *→* −*s*(*t*), *h →* 1 − *h*. For constant selection, it has been claimed that looking backward in time (that is, *t → T*_*c*_ − *t*), the wildtype frequency obeys the same dynamics as the mutant frequency with *h →* 1 − *h* but *s* unchanged (Martin and Lambert, 2015); however, this prescription results in a forward Kolmogorov equation with negative population size which is clearly absurd. Therefore, we will work always looking forward in time.

Proceeding in a manner similar to that for stochastic phase *A*, the distribution 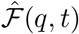 for the wildtype frequency subject to the initial condition *q*(*t*_2_) can be found for *t > t*_2_. Then the probability that the wildtype goes extinct by time *T*_*c*_ is given by (refer (15) for a comparison)

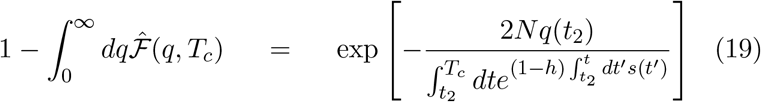

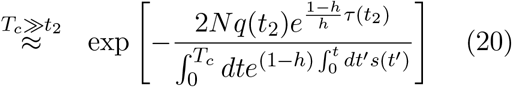

On taking the derivative of the above cumulative distribution with respect to *T*_*c*_ and averaging over the distribution of *q*(*t*_2_) which can be found using (16) and (18), we finally arrive at the distribution of conditional fixation time for a mutant with initial phase *θ*:

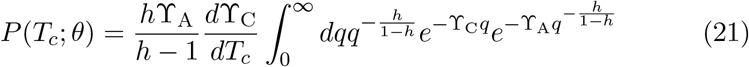

where

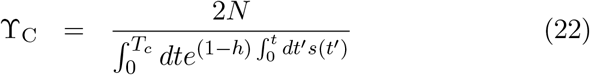

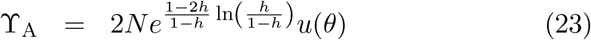

and the eventual fixation probability *u* is given by (15) for a single mutant.

Figure 4 shows a comparison between the expression (21) and the results obtained using numerical simulations when the cycling frequency is below, above and close to the resonance frequency *ω*_*r*_ (Devi and Jain, 2020), and we find a good agreement in all the three cases. Note that while the distribution is bell-shaped away from the resonance frequency, it is bimodal close to *ω*_*r*_ which results in a large variance in conditional fixation time (see inset of Fig. 4). For constant selection, we find that the generating function for the conditional fixation time obtained using (21) reduces to (A.11) of Martin and Lambert (2015).

**Figure 4:**
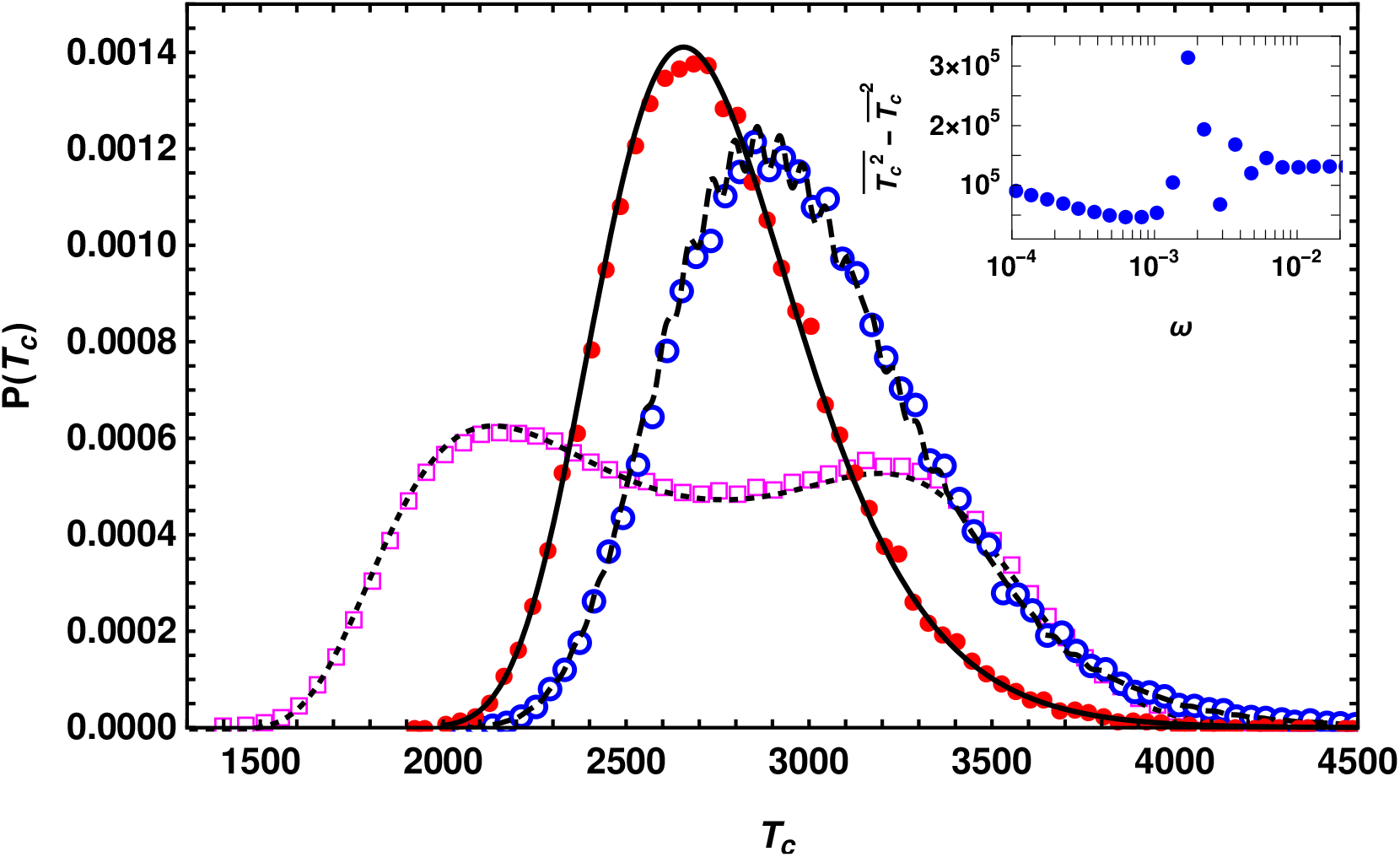
Fixation time distribution *P* (*T*_*c*_) when the mutant is beneficial at all times. The points and curves are obtained, respectively, from numerical simulations and the semi-deterministic result (21) for cycling frequency below, above and close to the resonance frequency *ω*_*r*_, and given by *ω* = 10^−4^ (•, solid line), 0.1 (○, dashed line) and 0.002 (◻, dotted line), respectively. The other parameters are *N* = 10^5^, *θ* = 0, 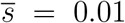, *σ* =0.007, *h* = 0.5. The inset shows that in accordance with the broad distribution close to *ω*_*r*_, the variance in the conditional fixation time is large in the vicinity of the resonance frequency.

### 4.4 Mean fixation time in slowly changing environments

Figure 5 shows that, except for strongly recessive or dominant mutant, the conditional mean fixation time 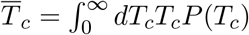 depends weakly on dominance for arbitrary rate of environmental change. As discussed in the last section, in slowly changing environments with increasing selection, 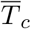 is smaller than the corresponding result in the static environment; in fast changing environments, it approaches the conditional mean fixation time in time-averaged environments and at intermediate frequencies, the fixation time 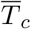 differs substantially from that in the constant environment.

**Figure 5:**
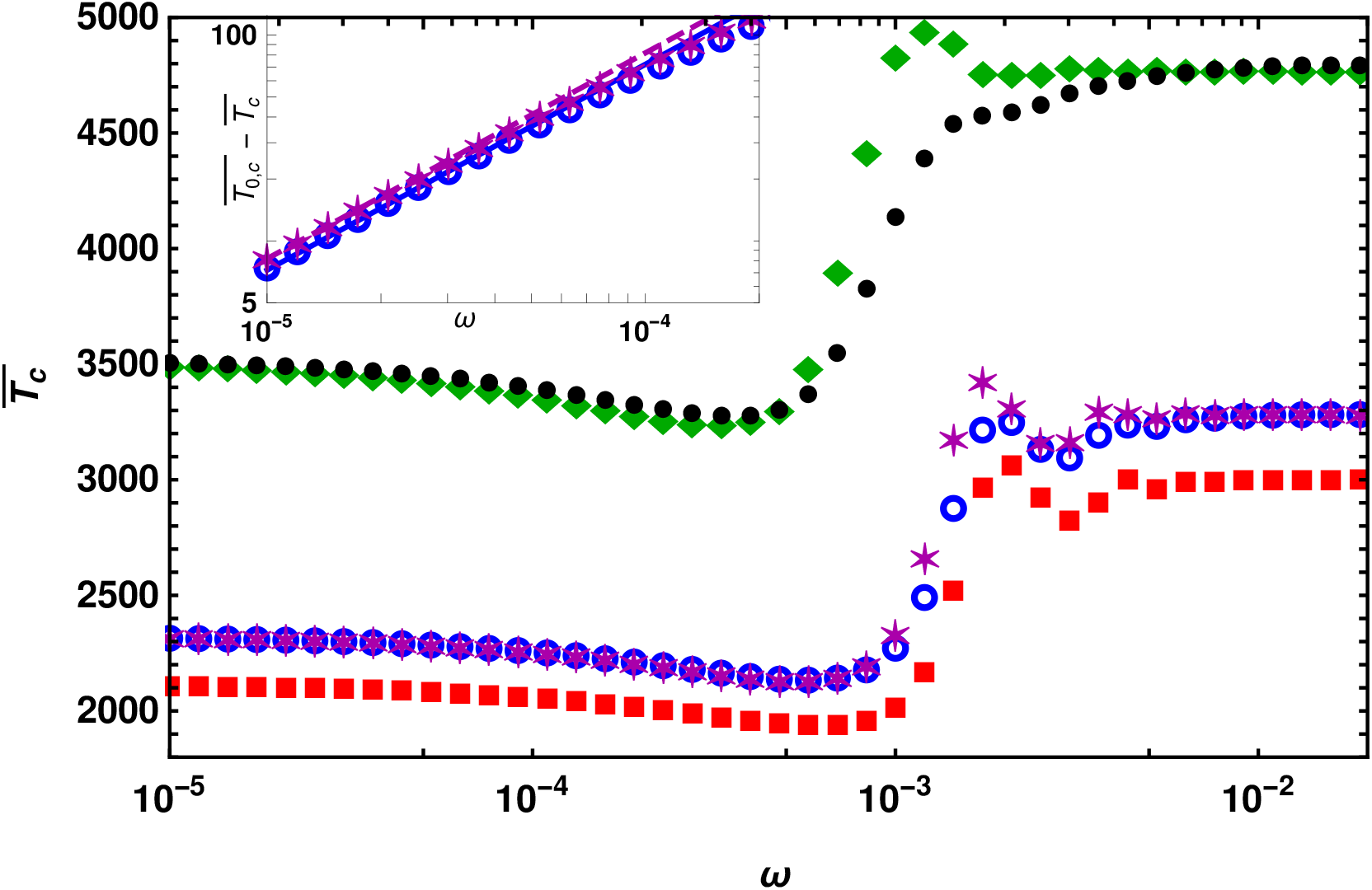
Conditional mean fixation time 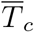 when the mutant is beneficial at all times to show that it depends weakly on the dominance coefficient. The points are obtained by numerically calculating (24) for *h* = 0.1 (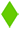), 0.3(*), 0.7 (○), 0.5 (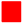), 0.9(•). The other parameters are *N* = 10^5^, 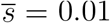, *σ* = 0.007, *θ* = *π/*4. The inset shows a comparison between (24) (points) and (26) (line) for the deviation in conditional mean fixation time in a slowly changing environment where *ω* ≪ *s*(0) ≈ 0.014.

Figure 5 also suggests that for small cycling frequencies, the conditional mean fixation time 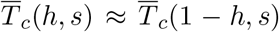. To understand this, using (21), we rewrite the time 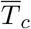 as

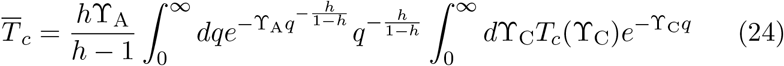

and analyze it for slowly changing environments using a perturbation theory. As explained in Appendix D, for *ω* ≪ *s*(0), we get 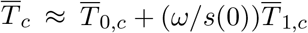 where,

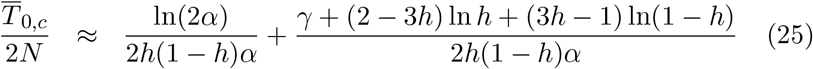

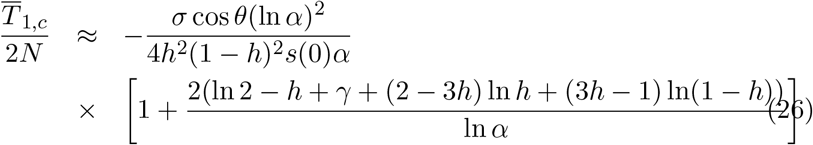

and, *γ* ≈ 0.577 is the Euler constant and as before, *α* = *Ns*(0). (Note that unlike here, in the last section, the fixation time 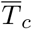 is expanded in powers of *Nω*.)

The semi-deterministic approach described here is not a systematic, controlled approximation (unlike various perturbation theories) and it is not clear how good this approximation is; however, here we find that (25) matches exactly with (4.2) of Ewing *et al*. (2011) (on replacing *N* in the above expression by 2*N*) which is obtained using a diffusion theory, and shows that the conditional mean fixation time in a population with dominance coefficient *h* is approximately equal to that in a population with corresponding parameter 1 − *h*. Note that this result holds for large *α*(≳10^3^) and for weaker selection, the dominant mutant takes longer than the recessive one to fix, as shown in Fig. 3 (see also Teshima and Przeworski (2006)).

Equation (26) captures the effect of slowly changing environment on conditional mean fixation time and matches well with the data obtained by numerically integrating (24) as shown in the inset of Fig. 5. The leading term on the right-hand side (RHS) of (26) is symmetric about *h* = 1*/*2 pointing to the approximate symmetry, 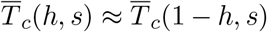 discussed above. However, the subleading correction does not have *h* ↔ 1 − *h* symmetry but its effect is small compared to the leading term for intermediate dominance. Figure S2 further suggests that the distribution of the fixation time has *h* ↔ 1−*h* approximate symmetry for small cycling frequencies but not for frequencies close to the resonance frequency in accordance with the behavior of the mean fixation time.

Equation (26) also shows that the mean fixation time decreases (increases) if the beneficial allele arises when the selection gradient (in time) is positive (negative), as intuitively expected; furthermore, both 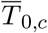 and 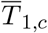 decay with increasing selection. Finally, we mention that at the beginning of this section, we had assumed that 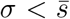. But in a slowly changing environment, the semi-deterministic approximation may be expected to work for 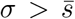 and 0 *< θ < π*; this is indeed confirmed in Fig. S2.

## 5 Data availability

The authors state that all data necessary for confirming the conclusions presented in the article are represented fully within the article.

## 6 Discussion

In this article, we have studied how an environment that is varying periodically and predictably in time affects the fixation time of a mutant in a finite, diploid population.

### Effect of the environmental parameters

We find that if the environment changes fast, the fixation time in the temporally varying environment differs considerably from that in the static environment, as can be seen in Figs. 1 and 5 at intermediate cycling frequencies. But for a meaningful comparison with the body of work on selective sweeps that assume constant environment (Stephan, 2016), here we have focused on the effect of slowly changing environments.

It should be noted that for time-dependent selection coefficient, the stochastic process is time-inhomogeneous and therefore the fixation time depends on the time at which the mutant arose (Uecker and Hermisson, 2011; Devi and Jain, 2020). For example, if the mutant arises when the selection is decreasing slowly with time towards the time-averaged selection coefficient 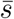 (as in Fig. 3), due to the reduced selection strength, the mutant takes longer to fix than if the selection stays constant at 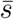.

### Selection regimes

In static environments, the qualitative behavior of the conditional mean fixation time of a mutant depends on the sign and strength of the scaled selection coefficient, *α* = *Ns*. For a beneficial mutant, if the selection is weak (0 *< α* ≪ 1), the fixation time increases with the dominance coefficient *h* and selection strength *α*, and can even exceed the fixation time for a neutral mutant (Mafessoni and Lachmann, 2015). But for moderately strong selection (1 ≪ *α* ≪ 100), the conditional mean fixation time decreases with *α* and increases with *h* (Teshima and Przeworski, 2006; Charlesworth, 2020), and for stronger selection, it decreases with *α* and is approximately same for two mutants with the same selection coefficient but dominance coefficients *h* and 1 − *h* (van Herwaarden and van der Wal, 2002; Ewing *et al*., 2011). The patterns for deleterious mutations follow on realizing that the conditional fixation time for a beneficial mutant with dominance coefficient *h* and deleterious mutant with same magnitude of selection but dominance level 1 − *h* are equal (Maruyama, 1974; Maruyama and Kimura, 1974).

Here, we find that all the qualitative patterns described above continue to hold if the environment changes slowly. However, there are quantitative differences: the exact symmetry between the conditional fixation time for beneficial and deleterious mutant mentioned above breaks in the changing environment. But, perhaps surprisingly, the approximate symmetry between the conditional fixation times for the beneficial mutants continues to hold. In fact, the conditional mean fixation time for a beneficial mutant is found to be mildly affected by the slowly changing environment but it is strongly impacted for a deleterious recessive mutant under moderate selection.

### Implications

In constant environment, due to the Maruyama-Kimura symmetry for conditional mean fixation times (Maruyama, 1974; Maruyama and Kimura, 1974), similar diversity patterns for beneficial and deleterious sweeps may be generated (Johri *et al*., 2020). But in changing environment, due to symmetry-breaking, a beneficial mutant in a mildly improving (deteriorating) environment can generate a diversity pattern different from that due to the fixation of a deleterious mutant under same selection pressure but in slowly deteriorating (improving) environment.

As already mentioned, although the qualitative patterns for fixation time in static environment are robust with respect to a slow change in the environment, there are quantitative differences. As a result, the effect of varying environment may be interpreted as an effective selection coefficient or dominance parameter. For example, in Fig. 3, the time of fixation for *h* = 1*/*2 and *α* = −14.14(*θ* = 5*π/*4) in slowly changing environment is about 1691. But if one assumes a constant environment, this fixation time is obtained for *α* = −15.27 which implies an 8% increase in the selection coefficient.

We therefore suggest to include the effect of changing environment in theoretical models of selective sweeps as this can potentially allow one to distinguish between the sign of selection and for correct estimation of the parameters.

### Open questions

Although this work allows us to delineate the parameter space where the slowly changing environment can potentially have a significant impact, we have not addressed precisely how the patterns of variability and other related statistics are affected by a changing environment and plan to study this in a future work. Generalizing the above results to include the effect of inbreeding and sex-linked inheritance (GlÉmin, 2012; Hartfield and Bataillon, 2020) could also help to assess the importance of changing environment in evolutionary dynamics.

## Acknowledgments

We thank Guillaume Martin and Amaury Lambert for helpful correspondence on their work.

## Appendix A Diffusion theory for time-inhomogeneous process

For a large finite population with small selection coefficient, the average fixation time can, in principle, be studied using the backward Fokker-Planck equation with time-dependent selection coefficient. The probability distribution Φ_b_(*x, t*|*p, t*_0_) that the mutant frequency is *x* at time *t*, given that it is *p* at time *t*_0_ *< t* obeys the following partial differential equation:

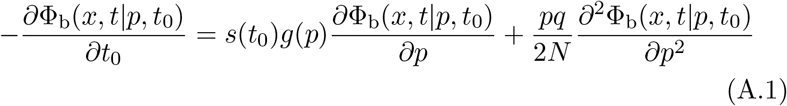

where 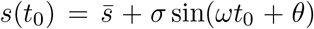 and *g*(*p*) = *pq*(*p* + *h*(1 − 2*p*)). In the above equation, the first term on the RHS is obtained on using that the deterministic rate of change of the mutant allele frequency is given by *dp/dt* = *s*(*t*)*g*(*p*), and the second term is due to the sampling noise in a finite population. Note that since the Fokker-Planck equations are derived by varying either the initial or the final time, not both (see, for e.g., Chapter 4, Risken (1996)), the backward Fokker-Planck equation (S5) in Uecker and Hermisson (2011) for the time-inhomogeneous process is incorrect as it contains an additional time derivative with respect to time *t* on the left-hand side (LHS).

The (unconditional) mean fixation time for a mutant arising at time *t*_0_ is then given by

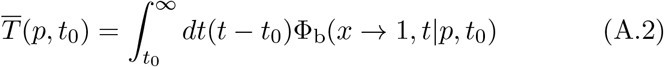

Using Leibniz integral rule,

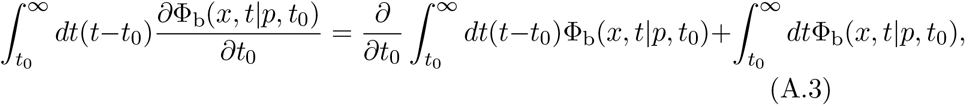

and (A.1), we then obtain

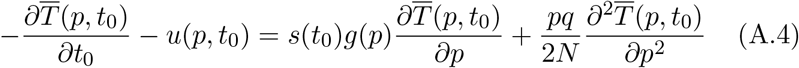

where the eventual fixation probability 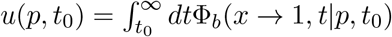 obeys (Uecker and Hermisson, 2011; Devi and Jain, 2020)

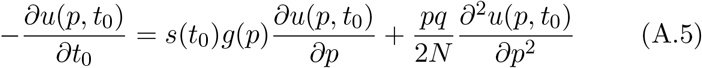

We verify that (A.4) and (A.5) reduce to the corresponding equations for the time-homogenous process where fixation probability and fixation time are independent of the initial time (Ewens, 2004). The partial differential equations (A.4) and (A.5) along with the boundary conditions *u*(0, *t*_0_) = 0, *u*(1, *t*_0_) = 1 and 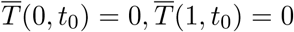 can, in principle, be used to find the mean fixation time for any sign of selection. However, these equations do not appear to be solvable, even for the dominance parameter *h* = 1*/*2, as the eigenfunction expansion method commonly employed for solving partial differential equations with time-dependent coefficients requires the eigenfunctions of the problem with constant selection that are, unfortunately, not known in a closed form (Jain and Devi, 2020).

## Appendix B Mean fixation time of on-average deleterious mutant

Below we consider the conditional mean fixation time of a co-dominant mutant in slowly changing environment (*ω* ≪ *σ* ≪ *N* ^−1^) which is neutral on average 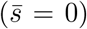 and can be written as 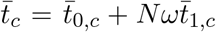 with 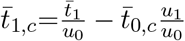 (see (7)). In a static environment with selection coefficient *s*(0) = *σ* sin *θ*, the eventual fixation probability *u*_0_ and the unconditional mean fixation time 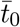 are given by (Ewens, 2004)

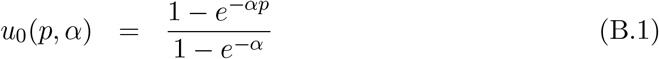

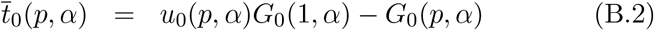

where

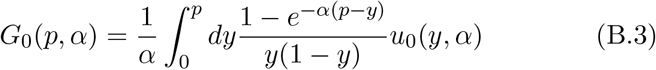

and *α* = *Nσ* sin *θ*. In a slowly changing environment, due to (3) and (4) in the main text, the corresponding quantities are given by

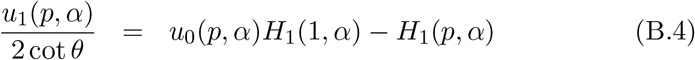

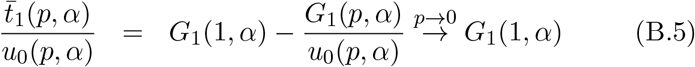

where

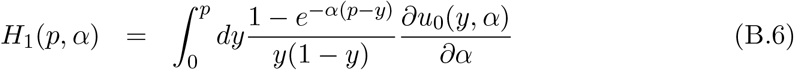

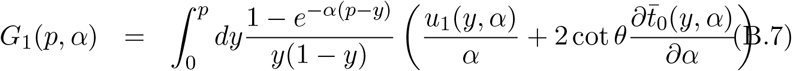

As discussed in the main text, for strong selection, the conditional mean fixation time for a mutant that remains beneficial until it fixes can be found within a semi-deterministic approximation and given by (26). Here, we therefore consider the difference 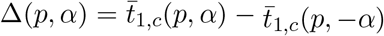 which, for single initial mutant, is given by

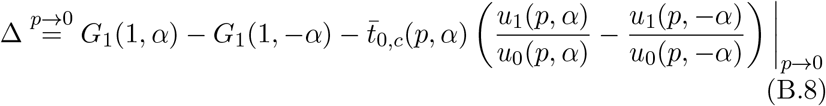

To find Δ, we require the conditional mean fixation time 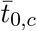 and the fixation probability *u*_1_ for arbitrary initial frequency. Although one can write exact expressions for them in terms of the exponential integrals, here we are interested in the strong selection regime (|*α*| *≫* 1). Using (5.1.10) and (5.1.51) in Abramowitz and Stegun (1964), to leading order in large *α*, we obtain

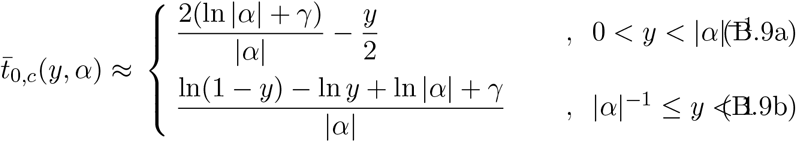

which decreases with the increasing frequency of initial mutants, as expected. For an initially beneficial mutant (that is, *s*(0) *>* 0), the fixation probability can be approximated by

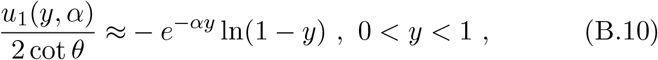

while for a mutant with *s*(0) *<* 0, we have

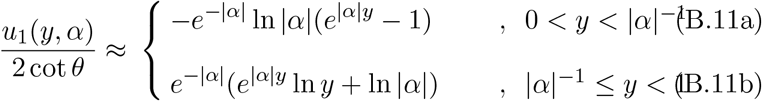

The above expressions (B.10) and (B.11a) reduce, respectively, to (11b) and (11c) of Devi and Jain (2020) when a single mutant is initially present.

Using (B.9a), (B.10) and (B.11a), we find that

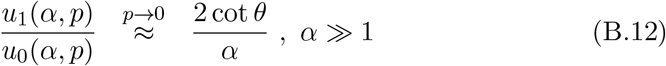

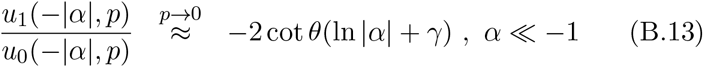

and therefore the last term on the RHS of (B.8) is given by 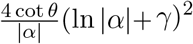. To estimate the difference, *G*_1_(1, *α*) − *G*_1_(1, −*α*), we first note that for a codominant mutant, the conditional fixation time, 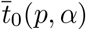 is symmetric about *α* = 0 for *any* initial frequency *p* (refer Sec. 5.4, Ewens (2004)). From the definition (B.7), we then get

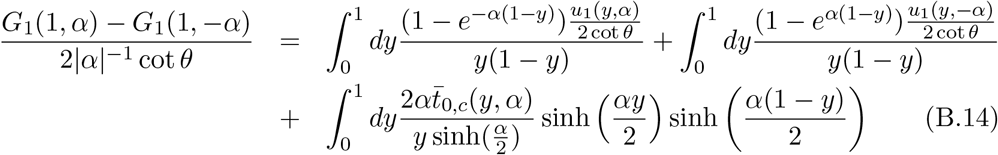

Using the results for 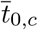 and *u*_1_ found above in the last equation and carrying out the integrals, we find that the first integral on the RHS is of order 1*/*|*α*|, the second integral, to order one, is equal to 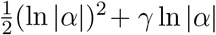 and the last integral is given by 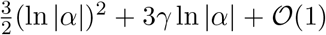. Putting all these results together shows that Δ decays as 1*/*|*α*| or faster. As the order one terms in (B.14) seem rather hard to calculate, we also studied these integrals numerically and find our numerical analyses to be consistent with Δ *∼* |*α*|^−1^. Thus, while 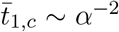 for beneficial mutant under strong selection (refer (26)), the change in the conditional mean fixation time for deleterious mutant decays slowly as |*α*|^−1^.

## Appendix C Feller process with time-dependent coefficients

Taking the Laplace transform on both sides of (12), we find that 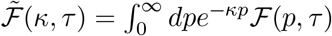 obeys a first order differential equation,

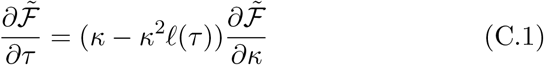

where 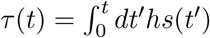 and 𝓁 (*τ*) = (2*Nhs*(*t*))^−1^. The above differential equation can be solved using the method of characteristics for the initial condition *ℱ* (*p*, 0) = *δ*(*p* − *p*_0_), and we obtain (Feller, 1951b; Masoliver, 2016)

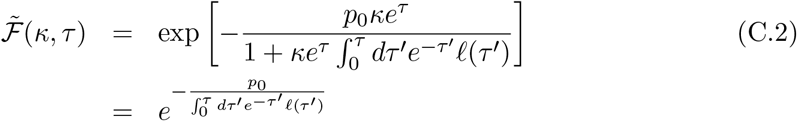

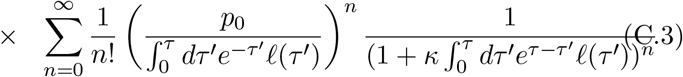

Taking the inverse Laplace transform of the summand in the last expression and then carrying out the sum over *n*, we get (13) in the main text. Note that the eventual fixation probability (15) can also be written as 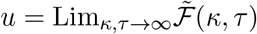.

## Appendix D Mean fixation time of on-average beneficial mutant

To find the conditional mean fixation time given by (24), we need to express *T*_*c*_ as a function of Υ_C_ using (22) which is given by

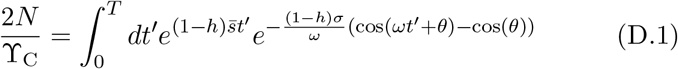

For small *ω*, we first expand the exponent of the integrand on the RHS to linear order in cycling frequency to obtain

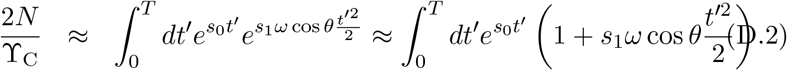

where *s*_0_ = (1−*h*)*s*(0) and *s*_1_ = (1−*h*)*σ*. On carrying out the integrals and taking the logarithm on both sides, to order *ω*, we get

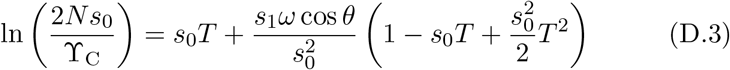

which can be inverted to finally give

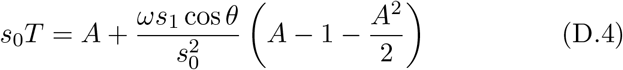

where 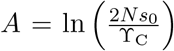 . Using this expression in the inner integral of (24) and carrying out the integrals, to leading and subleading orders in *α*, we obtain (25) and (26) in the main text.

## Supplementary Material

**Figure S1:**
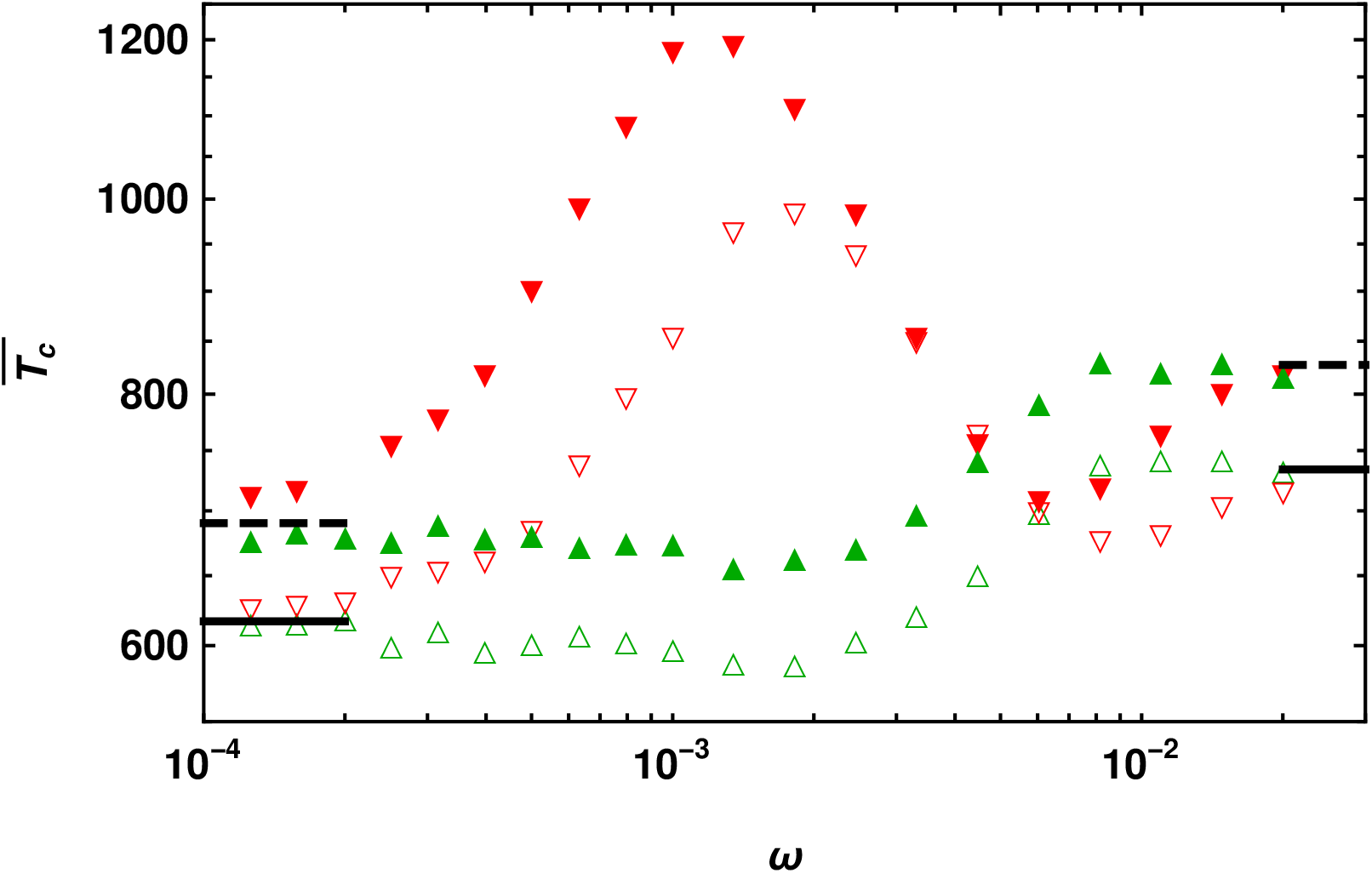
Conditional mean fixation time 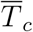 of a recessive (*h* = 0.3, 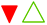) and dominant mutant (*h* = 0.7, 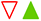) with selection coefficient 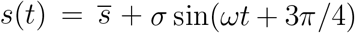 and −*s*(*t*), respectively, obtained by numerical simulations. The conditional fixation time in static environment with selection |*s*(0) | (left lines) and in the time-averaged environment with selection 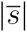 (right lines) are also shown for comparison, and obtained using diffusion theory for *h* = 0.3 (dashed) and 0.7 (solid). In all the cases, 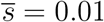, *σ* = 0.007 and *N* = 500.

**Figure S2:**
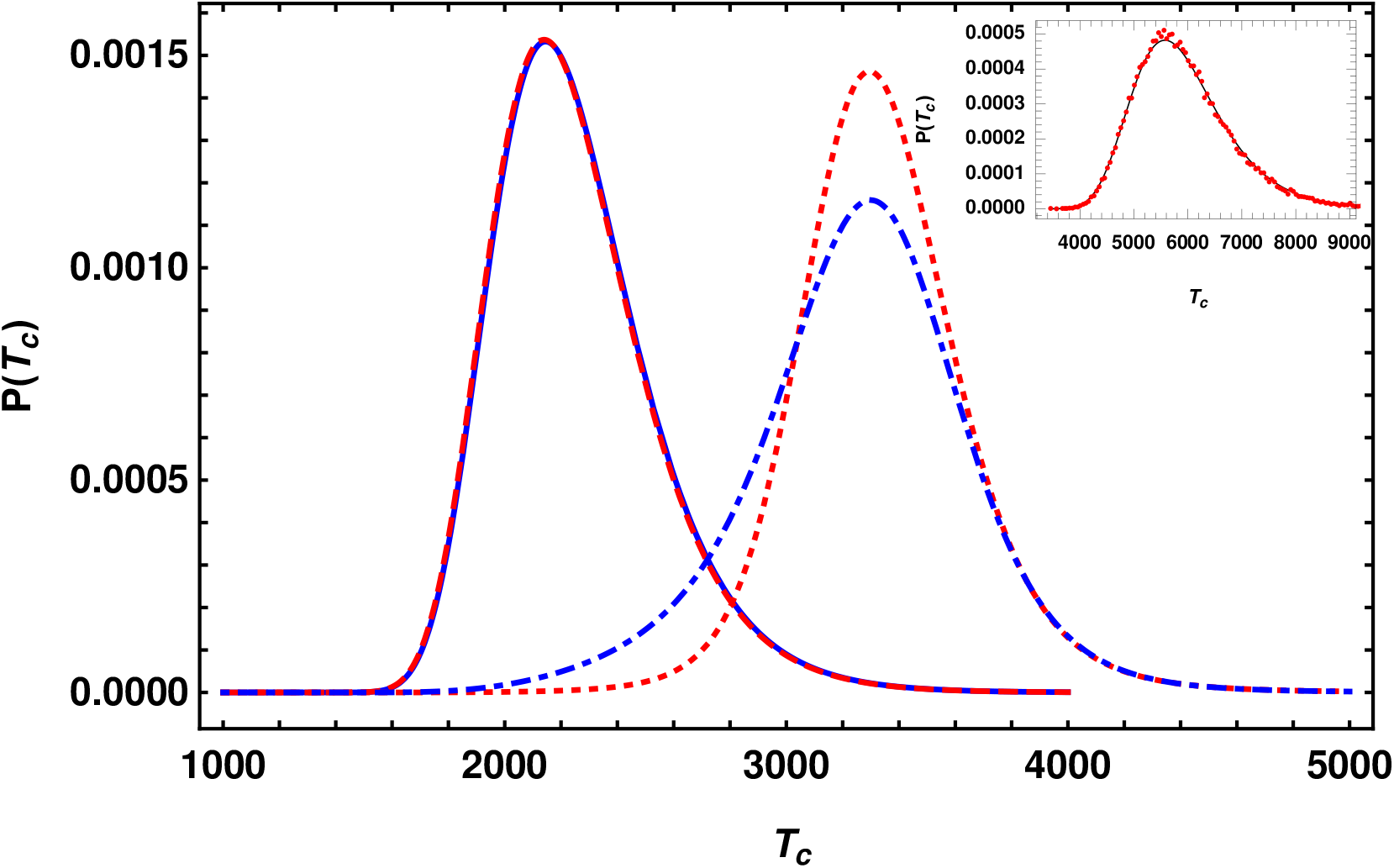
Distribution of conditional fixation time *T*_*c*_ obtained within semi-deterministic approximation and given by (21) when a mutant is beneficial at all times to show that it does not have the *h* ↔ 1 −*h* symmetry in fast changing environments. The left curves for *ω* = 10^−4^ show the distribution for *h* = 0.7 (solid blue) and 0.3 (dashed red) and the right curves for *ω* = 0.002 where *h* = 0.7 (dashdotted blue) and 0.3 (dotted red). The other parameters are *N* = 10^5^, *θ* = *π/*4, 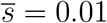, *σ* = 0.007. Inset: Distribution of fixation time of an initially beneficial mutant when time-averaged selection is zero and the environment changes slowly. The solid curve is obtained from (21) and the points are generated from simulations. The parameters are *N* = 10^5^, *σ* = 0.007, *ω* = 10^−6^, *h* = 0.3, *θ* = *π/*4.

